# ERK overstimulation leads to cell hyperproliferation in hereditary hemorrhagic telangiectasia landscape

**DOI:** 10.64898/2026.07.07.737030

**Authors:** José Luís Rocamora, Anna Casellas, Agnès Figueras, Pau Cerdà, Ferran Medina-Jover, Raquel Torres-Iglesias, Sandra D. Castillo, Mariona Graupera, Roxana Ola, Antoni Riera-Mestre, Francesc Viñals

## Abstract

Hereditary hemorrhagic telangiectasia (HHT) is a rare vascular disorder caused by pathogenic variants in members of the BMP9/ALK1 signaling hub. In the present study we show that, regardless of whether the alterations are caused by reduced BMP9/ALK1 signaling (pathogenic variants in the *ENG* or *ALK1* genes) or by overactivation of this pathway (such as the *SMAD6* pathogenic variants), all are associated with increased endothelial cell (EC) proliferation and high levels of ERK MAPK activation in patient biopsies. We reproduced this phenotype *in vitro* in ECs lacking SMAD6 or after SMAD1 knockdown using siRNA. Loss of SMAD6 leads to dysregulation of the Notch pathway, with downregulation of phosphatases and consequent overstimulation of ERK. In normal ECs, BMP9 and Notch signaling inhibit ERK activity by upregulating PPP1R3C, a regulatory subunit of the PP1 phosphatase. Notably, BMP9-mediated inhibition of ERK is abolished when cells are transfected with siRNA targeting PPP1R3C. ERK hyperactivation was also observed in an HHT2 mouse model (ALK1-2loxP;Cdh5-CreERT2). Loss of both ALK1 alleles in adult mice leads to vascular failure and hemorrhages in the lung and intestine; these injuries are significantly reduced by treatment with the MEK/ERK inhibitor selumetinib. Overall, our work identifies a key role for ERK activation involved in HHT pathogenesis, suggesting that ERK inhibition may represent a promising therapeutic strategy for these patients.

**Translational Perspective:** Hereditary hemorrhagic telangiectasia (HHTs) is a rare vascular disorder caused by mutations in members of the BMP9/ALK1 signaling hub. In the present study we show that all different forms of HHTs are associated with increased endothelial cell (EC) proliferation, which correlates with high levels of ERK activation in patient biopsies. ERK hyperactivation is also observed in an HHT2 mouse model in which loss of both ALK1 alleles in adult mice leads to vascular failure and hemorrhages in the lung and intestine. These injuries are significantly reduced by treatment with the MEK/ERK inhibitor selumetinib. Overall, our work identifies a key role for ERK activation in HHT pathogenesis, suggesting that ERK/MEK inhibitors may represent a promising therapeutic strategy for these patients.

## INTRODUCTION

Hereditary hemorrhagic telangiectasia (HHT) or Rendu–Osler–Weber syndrome is a rare autosomal dominant vascular disorder characterized by vascular malformations (VMs) in microvasculature, named telagiectases, and/or macrovasculature. Telangiectasis is the hallmark of HHT and consists in dilated postcapillary venules directly connected to dilated arterioles bypassing the normal capillary bed (Braverman *et al*, 1990). Telangiectases are typically found in the skin and mucous tissue, especially of the nose, mouth and lips, where they are more prone to hemorrhage causing epistaxis, gastrointestinal bleeding and anemia (Mora-Luján *et al*, 2020). Less frequently, vascular malformations (VMs) can appear in other organs, especially lung and liver, but also in the brain. Although HHT is inherited in a germline manner, its primary clinical manifestations do not typically present at birth but instead appear progressively over time. The disease exhibits age-dependent penetrance, with peak expression around the fourth and fifth decades of life except for brain arteriovenous malformations, which develop early in childhood, and for gastrointestinal involvement leading to anemia, which tends to worsen with age (Sánchez-Martínez *et al*, 2020).

HHT is caused by heterozygous loss-of-function variants in different genes coding for members of the BMP9/ALK1 signaling hub. ALK1 is a membrane type I receptor of the TGF-β receptor family predominantly expressed on endothelial cells (ECs). ALK1 specifically binds BMP9 and BMP10 with the cooperation of endoglin (ENG), a membrane co-receptor (Saito *et al*, 2017; Salmon *et al*, 2020). After ligand binding and activation, ALK1 intracellularly phosphorylates receptor-regulated SMADs (R-SMADs) SMAD1, 5 and 8. Phosphorylated R-SMADs form heteromeric complexes with SMAD4, they accumulate in the nucleus and regulate gene expression (Macias *et al*, 2015). The BMP9/ALK1 signaling pathway is essential for the regulation of angiogenesis, playing a key role promoting a more mature quiescent vessel phenotype and counteracting pro-mitogenic signals from factors such as VEGF or FGF-2 (Tillet & Bailly, 2014; Medina-Jover *et al*, 2022). There have been identified heterozygous loss-of-function variants in *ENG* (encoding endoglin protein) and *ACVRL1* (encoding ALK1 protein) genes in approximately 90% of cases referred for molecular diagnosis, leading to HHT1 (OMIM #187300) and HHT2 (OMIM #600376) respectively (McDonald *et al*, 2015). An additional 3% of cases present loss-of-function heterozygous variants in the *SMAD4* gene or in the in *GDF2* gene (encoding BMP9 protein). Recently, our group has identified a loss of function variant in *SMAD6* as a new driver in a family with an HHT-like phenotype (Cerdà *et al*, 2024). SMAD6 belongs to the inhibitory SMADs (I-SMAD) that downregulates BMP/SMAD signaling upon BMP9 stimulation (Miyazawa & Miyazono, 2017). Complete loss of SMAD6 results in vascular hemorrhage and embryonic lethality, emphasizing that balanced ALK1 signaling is crucial for ECs homeostasis (Galvin *et al*, 2000; Wylie *et al*, 2018). Disruption of this balance, either through excessive or insufficient ALK1 activity, can lead to vascular dysfunction and consequent HHT phenotype (Kulikauskas *et al*, 2023).

Consistent with its role in inducing quiescence by blocking proliferative signals, loss of function of the ALK1/BMP9 pathway causes cell cycle dysregulation and EC hyperproliferation in patients and animal models of HHT1 and HHT2 (Alsina-Sanchis *et al*, 2018; Iriarte *et al*, 2019; Genet *et al*, 2024). In this work we decided to study EC proliferative capacity in HHT-like disease caused by SMAD6 pathogenic variants, characterizing intracellular signaling pathways involved in its control.

## RESULTS

### Inactivating variant *SMAD6* p.Glu407Lys induces proliferation and ERK activation in ECs, as inactivacting variants in *ACVRL* do

We recently described that the *SMAD6* p.Glu407Lys variant had lost its inhibitory activity in the BMP/SMAD signaling pathway (Cerdà *et al*, 2024). As this variant has been identified in an HHT family as a novel genetic driver, we evaluated the impact of this mutation on the formation of vascular patterns by using hematoxylin-eosin and CD34 staining in a biopsy of a vascular malformation from a patient member of this family, and comparing it with telangiectasis biopsies from different HHT subtypes patients and a control skin biopsy from a non HHT patient (Fig. 1A and Sup. Fig. 1A). The vessels from the SMAD6 patient were of normal size but present in high density, although not as large as the vessels from classical HHT patients (Fig. 1A and Sup. Fig. 1A). We also studied the proliferative capacity of these vessels using Ki67 immunostaining. SMAD6 patient EC’s showed a higher proliferative capacity compared to ECs from normal skin vessels and to the proliferative capacity of ECs in different HHT forms samples (11.3% of ECs were Ki67 positive in the SMAD6 mutant compared to 2.5% in control ECs and 7.2% in the HHT patients, Fig. 1B).

**Figure 1.**
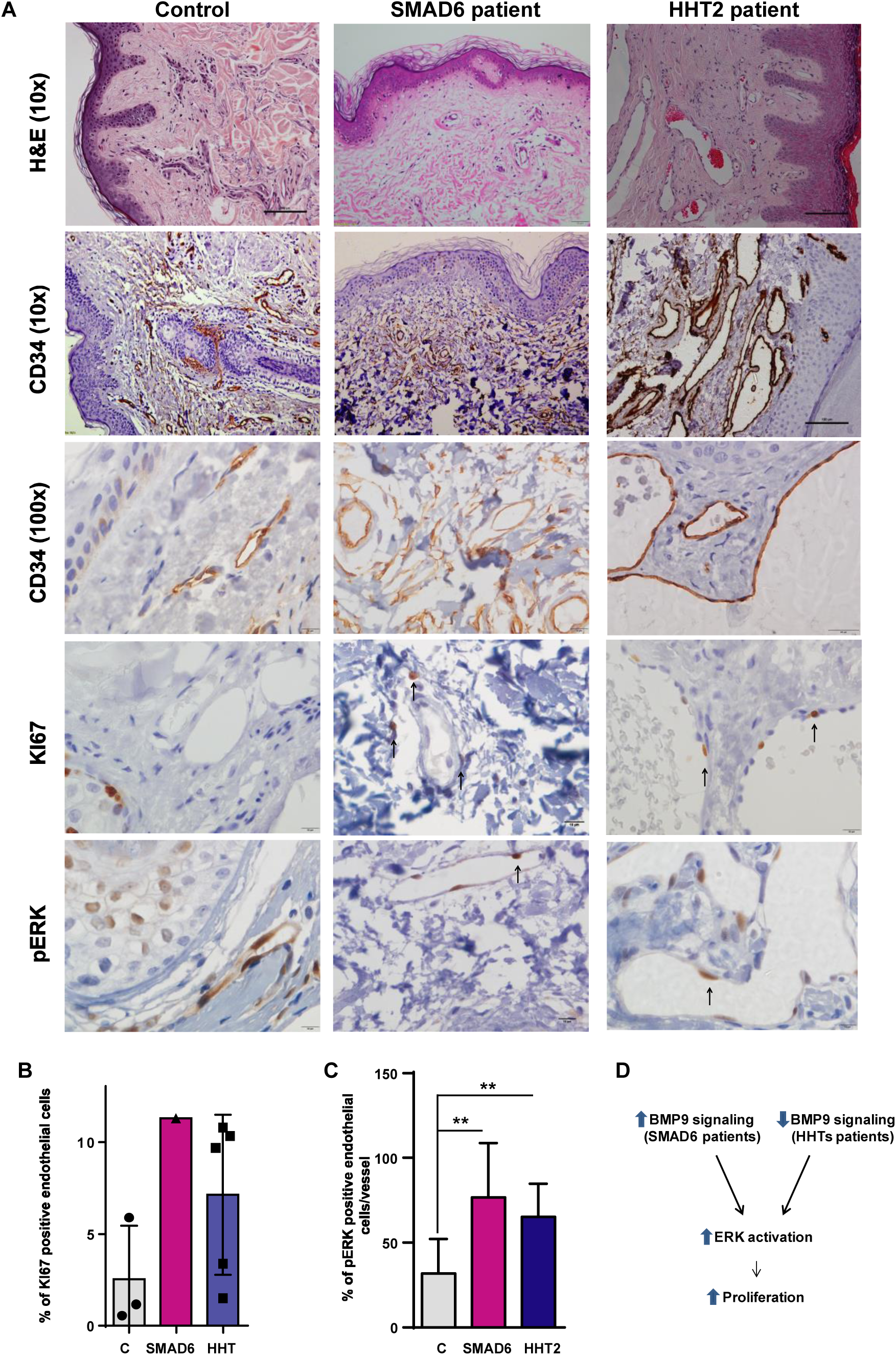
Different subtypes of HHT patient biopsies show inceased proliferation and ERK1/2 in vessel endothelial cells. **A)** Staining with a hematoxylin and eosin (x10), with anti-CD34 antibody (to stain vessels) (10x or 100x magnification), with anti-Ki67 (to stain proliferative cells) (x100) and with anti-pERK (x100) in control (left panel), *SMAD6* p.Glu407Lys patient (middle panel) and in a HHT2 patient (right panel) cross-sections of a paraffin-embedded human telangiectases. A representative image is shown. **B)** Quantification of the percentage of Ki67 positive ECs of the IHC shown in A) (represented Mean ± SD) (n=3 independent controls, n=1 *SMAD6* p.Glu407Lys patient and n=5 HHT1 or HHT2 independent patients). **C)** Quantification of the percentage of pERK positive nuclei relative to total nuclei in ECs of different blood vessels of the IF shown in Sup. Fig. 1B. Data are presented as mean ± SD (n = 10 independent pictures). Statistical significance was assessed using the Mann–Whitney U test. A p value < 0.05 was considered significant (** p<0.01). D) Model of data showed in Fig. 1A-C.

One of the key signaling pathways mediating EC proliferation is ERK MAPK (Srinivasan *et al*, 2009; Shin *et al*, 2016; Medina-Jover *et al*, 2020), so we analyzed if ERK was also stimulated in the patient with the *SMAD6* p.Glu407Lys variant or in those patients with classic HHT1 or 2. Therefore, we performed pERK immunohistochemistry on paraffin sections of the same samples previously analyzed. The results indicated that pERK was also increased in vessels from patients of both types of HHT compared to controls (Fig. 1A and Sup. Fig. 1A). This result was also obtained when immunofluorescence was performed, pERK, ENG (endothelial marker) and DAPI (nuclear marker) were stained (Fig. Sup. 1B), and pERK present in vessels was quantified (Fig. 1C). Therefore, there was an association between high proliferation and high pERK in HHT patients independently of the BMP9 signaling status (Fig. 1D).

### BMP9 signaling inhibits ERK activity by stimulating phosphatases in ECs

To identify at the molecular level how ERK was overactivated in these different HHT patients and, consequently, increase endothelial proliferation, we first studied the BMP9 hub in ECs in culture.

We started by modeling the HHT1 and HHT2 phenotype treating HUVEC with control siRNAs or siRNA against SMAD1, to reproduce the effect of lack of ALK1 or endoglin function (Fig. 2A and B). These cells were stimulated with VEGF and pERK was measured. As shown in Figure 2B, ERK was more stimulated under both basal and VEGF-stimulated conditions when SMAD1 levels were decreased by siRNA treatment, reproducing the phenotype observed in patients. We confirmed that BMP9 signaling negatively controls ERK activity by preincubating HUVEC 2-hour with BMP9 and then stimulating cells with VEGF for additional 30 min. BMP9 blocked basal and VEGF-mediated phosphorylation of ERK in HUVEC (Fig. 2C), without affecting total ERK protein levels as previously described (Alsina-Sanchis *et al*, 2018). In parallel, the phosphorylation and activation of upstream activators of the ERK pathway, such as MEK1 or Raf-1, were also repressed by the presence of BMP9 (Fig. 2C). In contrast, the ability of VEGF to stimulate its receptor VEGFR2 (Fig. 2C) or Ras (Sup. Fig. 2), the small G protein upstream of the Raf-MEK-ERK kinase cascade, was maintained in the presence of BMP9.

**Figure 2.**
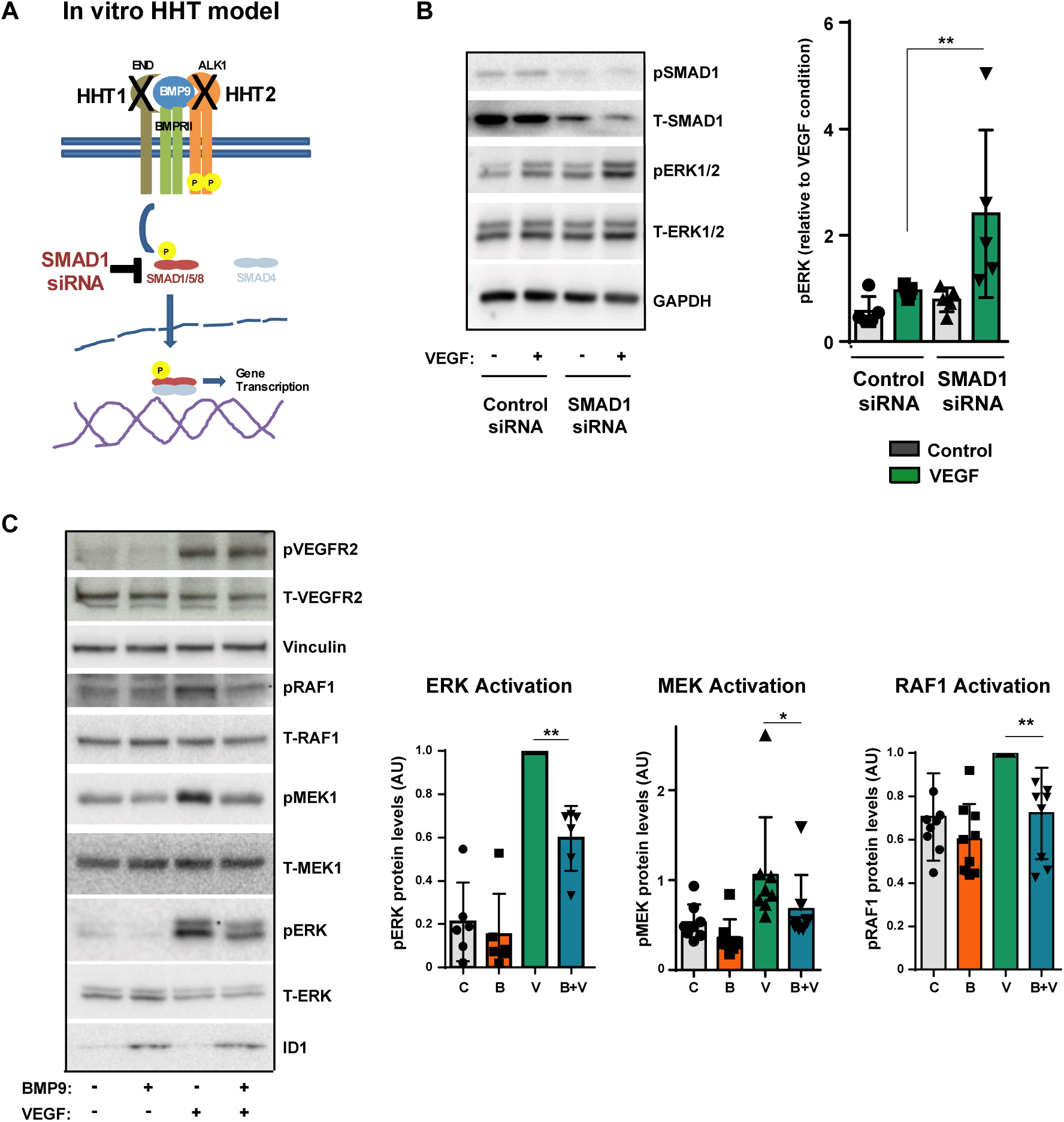
SMAD1 block induces ERK activation, while BMP9 treatment represses ERK. **A)** Model of HHT1 and 2 effects on BMP9 signaling. **B)** Left panel: Western blot assessing SMAD1 phosphorylation, total SMAD1 (T-SMAD1), ERK1/2 phosphorylation (pERK1/2), total ERK1/2 (T-ERK1/2) and GAPDH (as a loading control) in growth factor-depleted HUVECs stimulated (+VEGF) or not (-) with 10 ng/mL of VEGF for 15’. The two conditions were analyzed in the presence of a control siRNA or SMAD1 siRNA. A representative blot is shown. Right panel: Quantification of pERK normalized to VEGF-stimulated condition (n = 5 independent experiments). **C)** Left panel: Western blot assessing VEGFR2 phosphorylation (pVEGFR2), total VEGFR2 (T-VEGFR2), Raf-1 phosphorylation (pRaf1), total Raf1 (T-Raf1), MEK1 phosphorylation (pMEK1), total MEK1 (T-MEK1), ERK1/2 phosphorylation (pERK1/2), total ERK1/2 (T-ERK1/2), ID1 levels and Vinculin (as a loading control) in growth factor-depleted HUVECs pretreated with vehicle or 10 ng/mL of BMP9 for 2 hours, and with or without 10 ng/mL of VEGF for 15 additional minutes. A representative blot is shown. Right panels: Quantification of relative phosphorylation levels normalized to VEGF-stimulated condition of pERK (n= 6), pMEK1/2 (n= 9) or pRAF1 (n= 9 independent experiments).

Since VEGFR-Ras stimulation was not affected by BMP9 (Alsina et al. 2018 and present results), we hypothesized a possible negative regulation of the ERK pathway by one or more phosphatases activated by BMP9. In fact, other phosphatases already play a role in controlling BMP9 regulation in the case of the PI3K-AKT pathway, such as the phosphatase PTEN (Ola *et al*, 2016; Alsina-Sanchis *et al*, 2018). To validate this hypothesis, we first pretreated HUVEC with sodium orthovanadate, a general inhibitor of phosphatases. This reagent was able to completely block the negative effect of BMP9 on ERK phosphorylation in the absence of effects on BMP9 signaling (normal ID1 induction, Sup. Fig. 3A). Next, we decided to treat the ECs with okadaic acid, a specific inhibitor for members of the phosphoprotein phosphatases (PPPs) family (Silverstein *et al*, 2002), before their treatment with VEGF and/or BMP9. As expected, pretreatment with BMP9 clearly decreased pAKT and pERK under basal conditions (Fig. 3A). Furthermore, in the BMP9+VEGF condition, BMP9 was able to negatively affect VEGF-induced AKT and ERK phosphorylation. In the presence of okadaic acid, BMP9 signaling was not affected (ID1 induction) being still able to reduce pAKT levels in the BMP9+VEGF condition (Fig. 3A). In contrast, pretreatment with BMP9 failed to reduce pERK levels in the presence of okadaic acid. This result suggested that the okadaic acid-dependent phosphatases family were involved in the regulation of ERK by BMP9, but did not play a role in the regulation of AKT activation.

**Figure 3.**
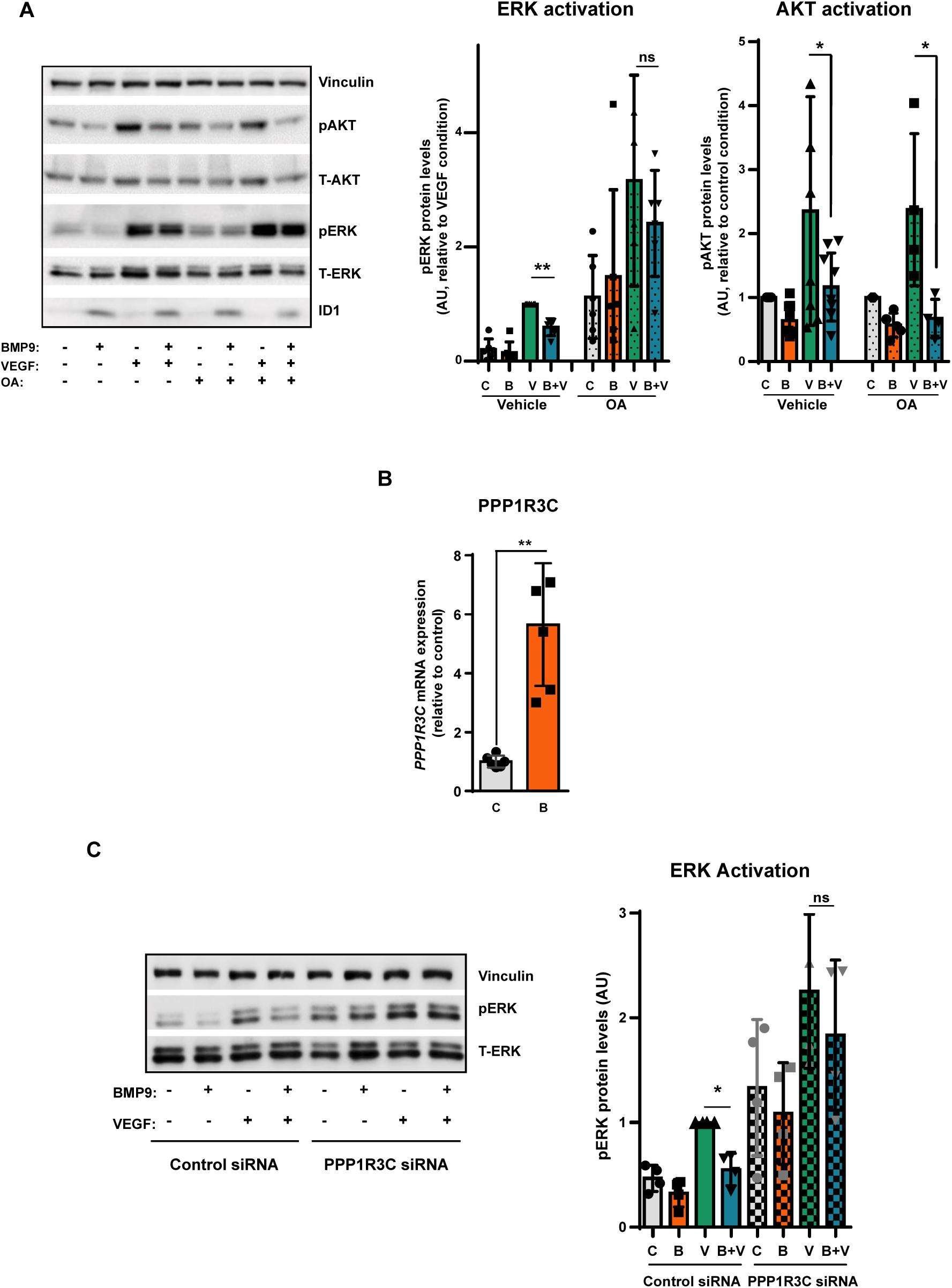
BMP9 controls ERK activation through activation of phosphatases. **A)** Left: Western blot assessing phosphorylated AKT (pAKT), total AKT (T-AKT), ERK1/2 phosphorylation (pERK1/2), total ERK1/2 (T-ERK1/2), ID1 levels and Vinculin (as a loading control) in HUVECs stimulated (+) or not (-) with 10 ng/mL of BMP9 (2h), VEGF (30’) or both. The four conditions were analyzed in the presence of a vehicle (DMSO) or 100 nM of okadaic acid (OA). A representative blot is shown. Right: Fold change of relative pERK (n=6 independent experiments, middle panel) or pAKT (n=6, right panel). Control (C), BMP9 (B) and BMP9+VEGF (B+V) were normalized to vehicle-treated VEGF (V) (pERK) or to control condition (pAKT). **B)** Fold change of *PPP1R3C* mRNA levels in HUVEC treated or not with 10 ng/ml BMP9 during 2 hours. Data are presented as mean ± SD (n= 6 independent experiments) normalized to control cells. **C)** Left panel: Western blot showing phosphorylated and total protein levels of ERK1/2 in HUVEC stimulated (+) or not (-) with 10 ng/mL of BMP9 for 2h, 10 ng/mL of VEGF for 30’ or both (n = 4 independent experiments). The four conditions were analyzed in the presence of a control siRNA or PPP1R3C siRNA. Vinculin was used as a protein loading control. A representative blot is shown. Right panel: Quantification of pERK normalized to the VEGF condition (n= 4 independent experiments). Data are presented as mean ± SD. Statistical significance was assessed using the Mann–Whitney U test. A p-value < 0.05 was considered statistically significant. (*p < 0.05; **p < 0.01).

Main targets of okadaic acid are the PPP family phosphatases PP1, PP2A and PP5. Accordingly, we sought to identify which of these phosphatases was/were regulated by BMP9. We treated HUVECs with BMP9 for 2 hours and measured mRNA or protein levels of several of the protein-forming parts (regulatory and catalytic) of the three-phosphatase complex expressed in the endothelium. Of the different subunits analyzed, only the regulatory subunit PPP1R3C of PP1 was significantly increased by BMP9 (Fig. 3B and Sup. Fig. 3B and C). To validate the involvement of this phosphatase in the repression of ERK activity induced by BMP9, we treated cells with control siRNAs or against PPP1R3C before stimulating them with BMP9 and/or VEGF. As shown in Fig. 3C, cells transfected with the siRNA against PPP1R3C increased basal levels of pERK, levels that were still stimulated by VEGF but in the absence of any effect of BMP9 on them. Taken together, these results confirmed that BMP9 controlled ERK activation in ECs through the expression and activity of phosphatases, such as PP1 and its regulatory subunit PPP1R3C, and that alterations in this signaling pathway (due to lack of functional BMP9 signaling as in patients with HHT1 and HHT2) could cause overactivation of ERK and of the proliferative capacity.

### *SMAD6^KO^* ECs exhibit increased proliferative capacity and ERK activity, despite higher BMP9/SMAD signaling

Next we went on to study what happened in the patient with a non functional SMAD6. To identify the mechanisms involved in the patient phenotype, we took advantage of the generation of immortalized SMAD6 knock-out (KO) dermal microvascular endothelial cells (hTERT-HDMEC) using CRISPR/Cas9 (Cerdà *et al*, 2024) (Fig. 4A). First, we confirmed that SMAD6 was increased in HDMEC control (Fig. 4B) and in HUVEC (Sup. Fig. 4A) by BMP9 treatment, probably as a negative feedback loop. *SMAD6^KO^* ECs exhibited higher BMP9/10signaling than control HDMEC cells by measuring SMAD1 phosphorylation after stimulation with BMP9 (Fig. 4B and Sup. Fig. 4B). The same result was obtained when BMP10 was used (Sup. Fig. 4C and D). Next, we measured how the absence of SMAD6 affected the proliferative capacity of these cells. Surprisingly, but confirming the results obtained in the patient samples, *SMAD6^KO^*ECs presented higher viability (Fig. 4C) and proliferation (measured by BrdU incorporation, Fig. 4D) than control HDMEC cells.

**Figure 4.**
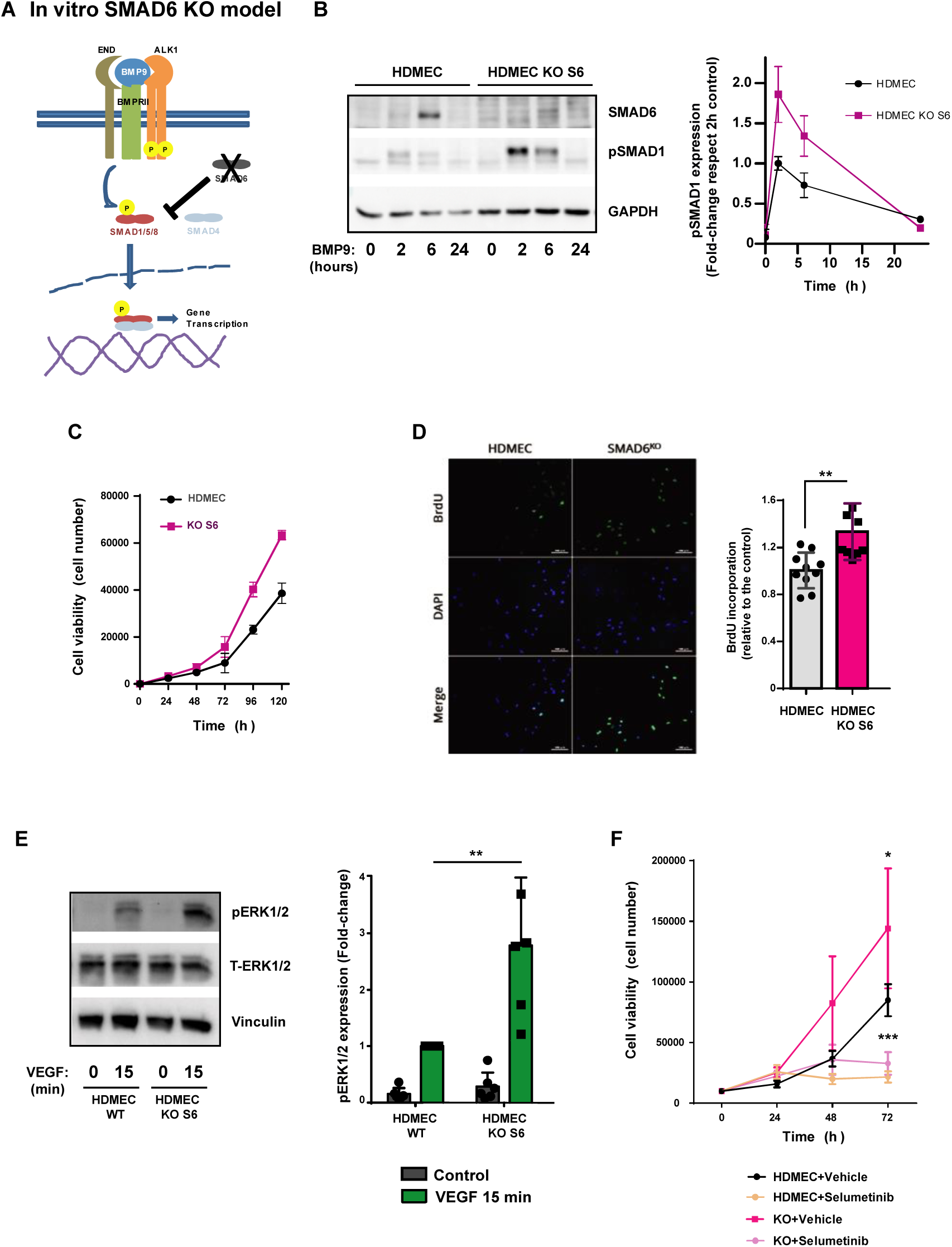
*SMAD6^KO^*ECs exhibit a higher BMP9/SMAD signaling but also a higher proliferative capacity. **A)** Model of lack of functional SMAD6 effects on BMP9 signaling. **B)** Left panel: Western blot assessing the SMAD6 expression, SMAD1 phosphorylation (pSMAD1) and GAPDH (as a loading control) in control (HDMEC) and *SMAD6^KO^* Ecs (HDMEC KO S6) treated with 1 ng/ml of BMP9 for 0h, 2h, 6h and 24h. A representative image is shown. Right panel: Quantification of pSMAD1 protein levels in the right (n=5 independent experiments). **C)** Number of control HDMEC or *SMAD6^KO^* ECs exponentially growing measured during five days (n=3 independent experiments). **D)** Proliferative capacity measured by BrdU incorporation in control HDMECs and *SMAD6^KO^* ECs (n=9 independent experiments) (right panel). Left panel: A representative image of BrdU staining. **E)** Left panel: Western blot assessing the ERK1/2 phosphorylation (pERK1/2), total ERK1/2 (T-ERK1/2) and Vinculin (as a loading control) in control HDMEC and *SMAD6^KO^*ECs treated with 10ng/ml of VEGF-A for 0 or 15 minutes. A representative image is shown. Right panel: quantification of pERK1/2 levels (n=6 independent experiments). **F)** Proliferation rates of control and *SMAD6^KO^*ECs treated with DMSO (–) or 3 µM selumetinib (+). Cell growth was monitored for 3 days, and values are expressed as mean ± SD from n = 6 independent experiments. Statistical significance was assessed using a two-way ANOVA with multiple comparisons against the control HDMEC (–) or control *SMAD6^KO^* ECs group, or the Mann–Whitney U test (**p < 0.005, ***p < 0.001).

We next analyzed the ability of VEGF to stimulate ERK in the absence of SMAD6. To do this, we depleted control HDMEC or *SMAD6^KO^*ECs and treated them or not for 15 minutes with VEGF-A. We then analyzed ERK stimulation by measuring the levels of phosphorylated and active ERK1/2. Surprisingly, the results indicated that ERK was more stimulated by VEGF in *SMAD6^KO^*ECs compared to control HDMECs (Fig. 4E). We decided to evaluate if this greater ability to stimulate ERK MAPK in *SMAD6^KO^* ECs influences their proliferative capacity. To this end, we inhibited MEK-1 (upstream of ERK) by treating cells with selumetinib, a clinically approved MEK inhibitor, and measured their proliferative capacity. Selumetinib was able to block the proliferative rate of both control and *SMAD6^KO^* ECs (Fig. 4F), confirming the involvement of ERK in controlling cell proliferation.

### Lack of SMAD6 decreases Notch signaling pathway and affects phosphatases expression

How can ERK be overstimulated in *SMAD6^KO^* ECs cells in a context of high BMP9/10 signaling? To study the molecular mechanisms involved in this alteration, we first compared the mRNA levels by RT-PCR of PPP1R3C, the PP1 subunit phosphatase that we had already identified as positively regulated by BMP9 and involved in ERK repression (Fig. 3B), between *SMAD6^KO^* and control HDMEC cells. *SMAD6^KO^*ECs showed an 80% drop in PPP1R3C mRNA levels compared to control cells (Fig. 5A), which could explain the over-stimulation of ERK in these cells. In order to identify the alterations caused by the absence of SMAD6 that could explain this dysregulation of PPP1R3C and consequently of ERK, we next performed an RNA sequencing analysis of exponentially growing *SMAD6^KO^* ECs compared to control HDMEC cells. Data clearly showed that the absence of SMAD6 caused a general change in RNA expression, as observed in the overall RNA expression analysis (Sup. Fig. 5A). When we studied the Top 10 Cell Regulation Biological Process affected, we confirmed differences in gene expression associated to cell proliferation, communication and activation cell processes among others (Sup. Fig. 5B). On the other hand, among all the genes identified in RNAseq altered in *SMAD6^KO^* ECs, different phosphatases stand out, including those subunits related to the control of PP1 activity that were also repressed, such as PPP1R1C, PPP1R3B and PPP1R3C (Sup. Fig. 5C), confirming the results obtained by RT-PCR. To validate that the observed effect of decreased of PPP1R3C in *SMAD6^KO^* ECs was due to the decrease in SMAD6 levels, we overexpressed the WT form of SMAD6 in *SMAD6^KO^*ECs and measured mRNA PPP1R3C levels. The results indicated that the observed decrease in PPP1R3C (from 2.55±0.08 to 1.46±0.17 log_2_ TPM) was recovered by 55% by overexpression of WT SMAD6 (from 1.46±0.17 to 2.06±0.36 log_2_ TPM).

**Figure 5.**
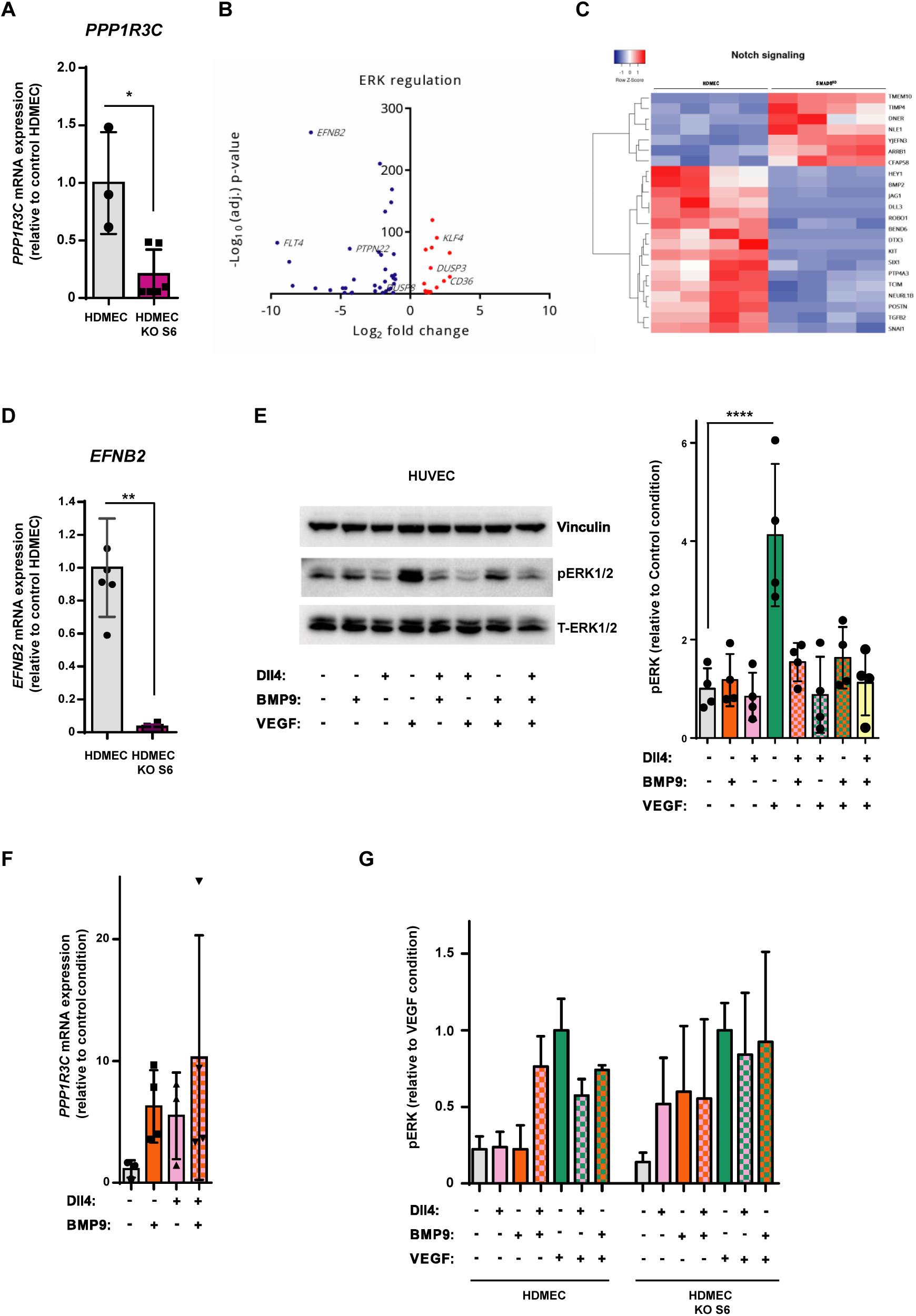
*SMAD6^KO^*ECs present alterations in Notch signaling and phosphatases gene expression. **A)** Fold change of *PPP1R3C* mRNA levels in *SMAD6^KO^*ECs relative to control HDMEC. Data are presented as mean ± SD (n= 6 independent experiments), normalized to control HDMEC. **B)** Volcano plot showing differentially expressed genes related to ERK regulation (GO:0070372) in *SMAD6^KO^* ECs compared to control HDMEC, based on RNA-seq analysis (n = 4 independent samples). **C)** Heatmap showing differentially expressed genes related to the NOTCH signaling pathway (GO:0007219) in *SMAD6^KO^*ECs compared to control HDMEC, based on RNA-seq data (n = 4). **D)** Fold change of *EFNB2* mRNA levels in *SMAD6^KO^*ECs relative to control HDMEC. Data are presented as mean ± SD (n= 6 independent experiments), normalized to control HDMEC. **E)** Western blot (left) assessing the ERK1/2 phosphorylation (pERK1/2), total ERK1/2 (T-ERK1/2) and Vinculin (as a loading control) in HUVEC treated or not (control) with 10 μg/ml coating Dll-4 for 24h, 10 ng/ml BMP9 for 2h or the combination of both in the presence or the absence of 10 ng/ml of VEGF-A for 15 minutes. A representative image is shown. Right: Quantification of pERK relative to total ERK in the control condition (n=4 independent experiments). **F)** Fold change of *PPP1R3C* mRNA levels in HUVEC treated or not with 10 μg/ml coating Dll-4 for 24h, 10 ng/ml BMP9 for 2h or the combination of both. Data are presented as mean ± SD (n= 4 independent experiments), normalized to control non treated HUVEC. **G)** Western blot assessing the ERK1/2 phosphorylation in *SMAD6^KO^* ECs relative to control HDMEC treated or not with 10 μg/ml coating Dll-4 for 24h, 10 ng/ml BMP9 for 2h or the combination of both in the presence or the absence of 10 ng/ml of VEGF-A for 15 minutes. The quantification of pERK is shown relative to the VEGF condition (n=3 independent experiments). Statistical significance was determined using the Mann–Whitney U test. A p value < 0.05 was considered statistically significant (* p<0.05, ** p<0.01, **** p<0.001).

To identify which signaling pathway altered in *SMAD6^KO^* ECs could generate this effect on PPP1R3C and ERK control, we also took advantage of RNA-seq data. We analyzed GEO “ERK regulation” in these cells compared to control HDMEC cells (Fig. 5B). Ephrin B2 (*EFNB2*) was one of the most downregulated genes in *SMAD6^KO^* ECs. This protein is a membrane receptor involved in Notch signaling, a pathway that has been implicated in the regulation of ERK activity during sprouting angiogenesis (Pontes-Quero *et al*, 2019). Cells lacking SMAD6 presented the entire Notch pathway repressed (Fig. 5C). We also confirmed low levels of EFNB2 mRNA by RT-PCR in independent samples of *SMAD6^KO^*ECs (Fig. 5D). Considering that Notch and BMP9 converge in the regulation of different genes that control ECs behavior (Larrivée *et al*, 2012; Kerr *et al*, 2015), we decided to examine the effect of both pathways, Notch and BMP9, on the activation of ERK by VEGF. First, we checked the effect of Dll4 (inducing Notch signaling), BMP9 or the combination on the mRNA expression of different downstream effectors of Notch signaling, such as the transcription factors Hey1 and Hey2, or the ligand Jagged1. Results indicated that BMP9 stimulated the production of these Notch targets in endothelial cells in a similar way to Dll4, without presenting synergistic effects (Sup. Fig. 5D). Pretreatment of control HUVEC with Dll4 decreased the ability of VEGF to stimulate ERK, even better that BMP9 did (Fig. 5E). When we pretreated cells with both signals, the effect was not additive, inhibiting ERK phosphorylation to the same extent as each factor alone. As expected, Dll4 also stimulated PPP1R3C mRNA in HUVEC as BMP9 did (Fig. 5F). Finally, when we evaluated the capacity of BMP9 or Dll4 to affect ERK stimulation by VEGF in *SMAD6^KO^*ECs compared with control HDMEC, the effect of both factors was decreased in the absence of SMAD6 (Fig. 5G).

Thus, all these results confirm that the absence of a functional SMAD6 causes a global shift in ECs behaviour, altering amongst others, the Notch pathway. This change in the signaling, together with the observed simultaneous overactivation of the BMP9 pathway, would generate modifications in phosphatases that regulate ERK and which would ultimately give rise to overactivation of ERK and greater proliferation.

### An animal model of HHT2 shows an increased ERK activation and blocking its activity paliates HHT injuries

Finally, to demonstrate the importance of ERK activity in HHT, we decided to use an adult HHT2 mouse model in which, to accelerate the onset of the disease, both *Acvrl1* alleles were specifically abrogated in ECs (*Acvrl1^EC-KO^*). To do this, we used the Cdh5-CreERT2;*Acvrl1^flox/flox^* mouse line (Sung *et al*, 2009; Ouarné *et al*, 2024) in which a Cre-ERT2 gene is under the control of the endothelium-specific Cdh5 promoter, allowing endothelium-specific deletion of *Acvrl1* after tamoxifen administration. *Acvrl1^flox/flox^* mice without Cre-ERT2 allele were used as control. Thus, 1mg of tamoxifen was administered intraperitoneally for five consecutive days to two-months-old mice, which resulted in Cre activation and deletion of both loxP-flanked *Acvrl1* alleles in ECs, generating *Acvrl1^EC-KO^* mice. These mice typically died 8–12 days after the first dose of tamoxifen due to severe gastrointestinal (Fig. 6A and B) and pulmonary (Fig. 6F and G) hemorrhage, with disorganization of intestinal crypts in the duodenum and ileum (Fig. 6B). Loss of BMP9 signaling in the *Acvrl1^EC-KO^* mice caused an increase in ERK phosphorylation in ECs of vessels in both the intestine (Fig. 6C-E) and lung (Sup. Fig. 6A). To assess whether inhibition of the ERK pathway could alleviate these symptoms, selumetinib (MEK inhibitor) (37.5 mg/kg) was administered for 8 days starting on the first day of tamoxifen injection. The experiment was terminated by sacrifice of the mice on day 8. The effect of selumetinib on ERK activation in blood vessels was observed by immunohistochemistry of pERK in treated intestines (Figure 6C-E) and lungs (Sup. Fig. 6A).

**Figure 6.**
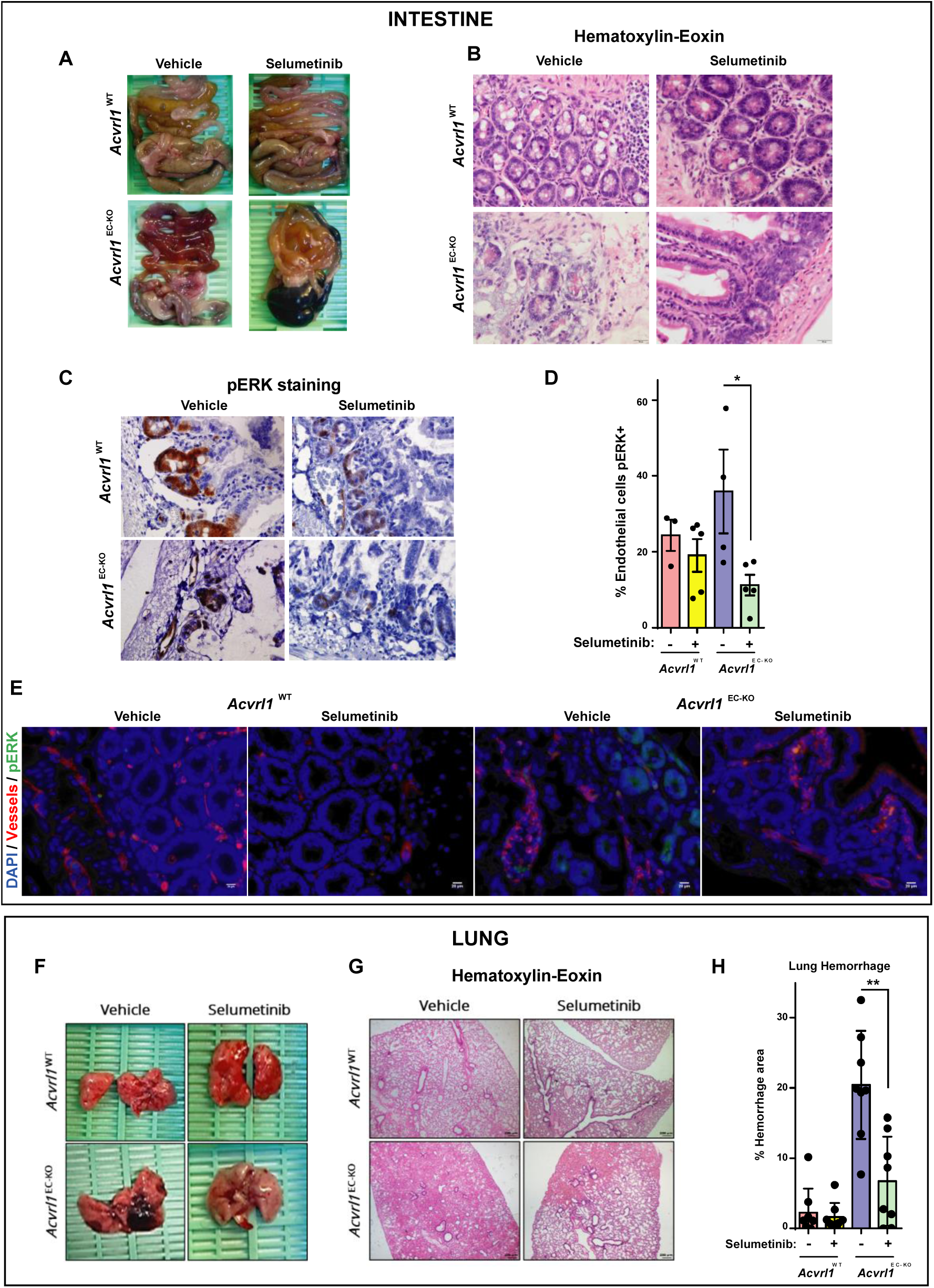
Selumetinib partially reverst hemhorragies in lungs and guts in a mouse HHT2 model. **A)** Macroscopic images of gut from *Acvrl1^flox/flox^ (Acvrl^WT^)* and Cdh5-CreERT2;*Acvrl1^EC-KO^* mice treated with vehicle or selumetinib. A representative image is shown. **B)** Hematoxylin-Eosin staining of sections of paraffin-embedded jejunums of gut from the same experimental groups. Images were captured at 10x magnification. A representative image is shown. **C)** pERK immunohistochemistry staining in sections of paraffin-embedded gut from the same experimental groups. A representative image is shown. **D)** Quantification of pERK positive endothelial cells by vessel in samples showed in C (n=5 independent animals). **E)** Immunofluorescence staining of pERK (green), vessels (red) and DAPI (blue) in cross-sections of paraffin-embedded gut from *Acvrl1^flox/flox^ (Acvrl^WT^)* and Cdh5-CreERT2;*Acvrl1^EC-KO^* mice treated with vehicle or selumetinib. A representative image is shown. **F)** Macroscopic images of lungs from *Acvrl1^flox/flox^ (Acvrl^WT^)* and Cdh5-CreERT2;*Acvrl1^EC-KO^*mice treated with vehicle or selumetinib. A representative image is shown. **G)** Hematoxylin-Eosin staining of sections of paraffin-embedded lungs from the same experimental groups. Images were captured at 10x magnification. A representative image is shown. **H)** % of hemorrhagic area in lungs (n = 8 independent animals) from *Acvrl1^flox/flox^ (Acvrl^WT^)* and Cdh5-CreERT2;*Acvrl1^EC-KO^*mice treated with vehicle or selumetinib, detected by iron present outside blood vessels. Statistical significance was assessed using the Mann–Whitney U test. A p value < 0.05 was considered statistically significant (*p<0.05, **p<0.01).

Treatment with selumetinib improved the structural integrity of intestinal crypts (Fig. 6B, right panels) and lungs (Fig. 6G, right panels) and significantly reduced pulmonary hemorrhage in *Acvrl1^EC-KO^* (Figure 6H).

## DISCUSSION

In this study, we demonstrate that all forms of HHT, regardless of the underlying causative variants, exhibit high levels of EC proliferation leading to abnormal vessel growth. Moreover, we identify ERK overactivation, driven by decreased phosphatase activity, as the common signaling alteration underlying this phenotype. Inhibition of ERK by treatment with the MEK/ERK inhibitor selumetinib in a HHT2 model in the adult significantly reduced the injuries and hemorrhages present in lung and intestine (Fig. 7).

**Figure 7.**
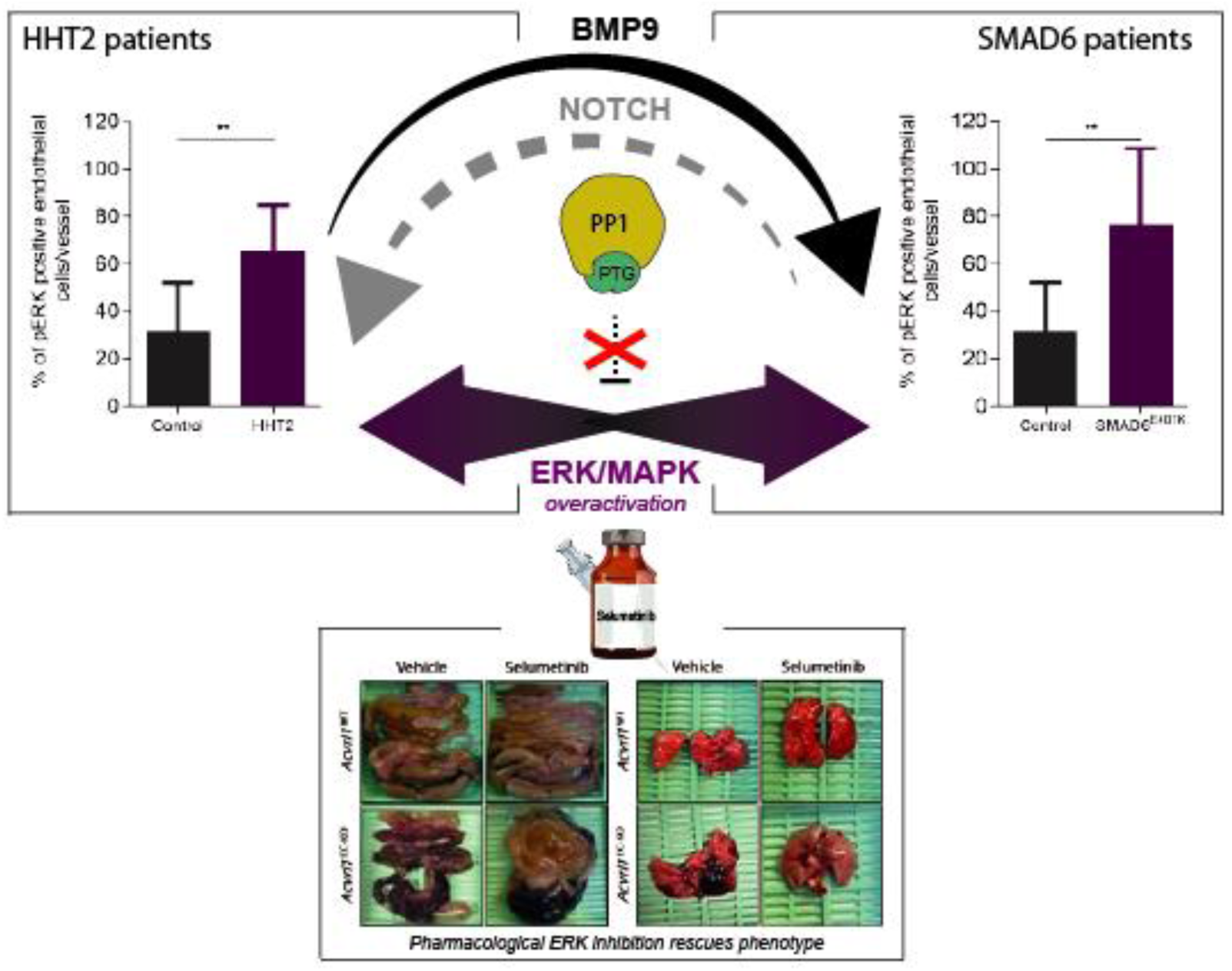
Model proposed. All forms of HHT, regardless of the underlying causative variants, exhibit low levels of phosphase PP1 and high levels of pERK. As a consequence, all present high endothelial cell proliferation leading to abnormal vessel growth. Inhibition of ERK by treatment with the MEK/ERK inhibitor selumetinib in a HHT2 model in the adult significantly reduced the injuries and hemorrhages present in lung and intestine.

Our data indicate that PP1 phosphatase is a key enzyme mediating ERK pathway inhibition downstream of BMP9 or Notch signaling. Protein phosphorylation is one of the most relevant post-translational modifications regulating most intracellular signaling pathways involved in cell behavior. Phosphorylation is a highly dynamic and reversible process, with most phosphorylated residues displaying half-lives of only seconds. The enzymes responsible for removing phosphate groups from proteins are the phosphatases. They play a crucial role controlling intracellular signaling pathways by terminating kinase cascades activated by a growth factor or stimulus, as the family of MAPK cascades (ERKs, JNKs and p38s), or by directly blocking kinase basal activity (Gelens *et al*, 2018). Several phosphatases have been implicated in the regulation of the ERK MAPK pathway (Raman *et al*, 2007), including dual tyrosine/threonine phosphatases (Owens & Keyse, 2007) and members of the phosphoprotein phosphatases (PPPs) family (Silverstein *et al*, 2002). PPPs are holoenzymes (multisubunit complexes) composed of catalytic and regulatory subunits that determine substrate specificity and catalytic efficiency (Shi, 2009; Brautigan & Shenolikar, 2018; Scheinost & Köhn, 2025). In this work we identified the regulatory subunit of PP1, PTG (“protein targeting to glycogen”), encoded by *PPP1R3C* gene, as a transcriptional target of BMP9 in ECs and involved in modulating ERK activity. PTG is the most ubiquitously expressed isoform among PP1 regulatory subunits (Printen *et al*, 1997; Semrau *et al*, 2022) and it is known to play a central role in glycogen metabolism. In ECs, ERK inhibition mechanisms have also been described in response to mechanical stress, in a tyrosine phosphatase-dependent manner (Shi *et al*, 2007), or upon contact inhibition (Viñals & Pouysségur, 1999), indicating that various phosphatases can repress ERK activity in response to distinct cytokines or cellular contexts. Our results suggest that the inhibitory effects of Notch (Pontes-Quero *et al*, 2019) and BMP9 (Medina-Jover *et al*, 2020) on VEGF-induced endothelial cell proliferation may be mediated by PP1 activation, leading to ERK pathway repression.

The second key finding of our study is that different forms of HHT share a common mechanism to increase endothelial proliferation, driven by ERK hyperactivation. Endothelial proliferation has been implicated in the development of HHT (Genet *et al*, 2024), as the dependence of endothelial proliferation on ERK signaling is well stablished (Srinivasan *et al*, 2009; Shin *et al*, 2016; Medina-Jover *et al*, 2020). Here we connect these concepts, demonstrating that proliferative ECs within HHT lesions display elevated ERK activity, regardless of the malfunctioning member of the BMP9/ALK1 signaling hub. According to the current model of HHT pathogenesis, local vascular malformations arise following a second hit such as inflamatory or angiogenic stress in a local zone of heterozygous individuals. Our results indicate that either, a lack of BMP9 signaling (in HHT1 and 2), or an excess of BMP9 signaling (SMAD6 deficiency) that completely alters the behavior of endothelial cells blocking Notch among other signals, both BMP9 alterations converge in impaired PP1 activity. This leads to higher basal activity of ERK or a greater capacity to stimulate this signaling pathway by proangiogenic factors such as VEGF and, in consequence, vascular malformations. Indeed, RNA-seq data analyses of blood outgrowth endothelial cells (BOECs) from HHT1 and HHT2 patients (Shovlin *et al*, 2024) revealed reduced expression of several PP1 subunits (PPP1R15, PPP1CA, PPP1CB, and PPP1CC), supporting the hypothesis that PP1 dysfunction contributes to ERK pathway overactivation in HHT and validating our cellular findings.

HHT treatments have traditionally relied on complications management and no therapies to treat HHT are licensed. The phenotypic heterogeneity in HHT leads to different clusters, including patients with predominantly severe epistaxis, transfusion-dependent GI bleeding, or pulmonary or hepatic predominant disease. Thus, new systemic therapies are needed to offer a more individualized and biologically based management, according to different clusters of HHT patients. Moreover, alternative drugs are also imperative for those patients with no response to a target therapy or if tachyphylaxis phenomena (sudden and progressive decrease in response after repetitive administration of a drug) occur (Villanueva *et al*, 2023). Our results point to a shared dependence on ERK signaling among HHT, proving that pharmacological ERK inhibition with selumetinib reduces proliferation and partially normalizes vessels in our HHT mouse model. Consistent with this, a recent case report (Shovlin *et al*, 2024) described an HHT patient treated with trametinib (a different MEK-inhibitor) for cancer who exhibited a marked reduction in bleeding episodes as a side effect. Altogether, highlights MEK-inhibitors as promising candidates for drug repurposing to treat HHT patients or developing a new specific drugs using ERK inhibition, as it occurred with engasertib, a new AKT inhibitor (Al-Samkari *et al*, 2025). Hence, the rellevance of our study providing the rationale for this new potential family drugs. Further investigation is needed to determine whether concomitant use of these new drugs (as PI3K, AKT or ERK/MAPK inhibition) provides additional benefit in the treatment of HHT patients.

## METHODS

### Patient samples

We selected patients attended in the HHT Unit in the Hospital Universitari de Bellvitge with a definite HHT diagnosis according to the Curaçao Criteria (meet ≥3 criteria) and a positive genetic test identifying variants in *ACVRL1*. The patient presenting the loss of function *SMAD6* p. (Glu407Lys) variant has been already described (Cerdà *et al*, 2024). A biopsy of a cutaneous thoracic VM from a patient member of this SMAD6 family and a punch biopsy (3 mm) from a cutaneous telangiectasis on the fingertip from an HHT2 patient were obtained under the usual conditions of sterility and hygiene by a senior dermatologist. Samples were encrypted according to a code assigned to each patient. According to Helsinki Declaration, all participants provided their signed informed consent for participation in the study following local Ethics Committee requirements. Accordingly, all individual-level data was de-identified. The study was approved by the Clinical Research Ethics Committee of the Hospital Universitari de Bellvitge (approval number PR329/23).

### Animal Procedures

*In vivo* experiments were carried out in agreement with the legislations and guidelines of the Catalan Ministry of Agriculture, Livestock, Fisheries and Food (Catalonia, Spain). Protocols were approved by the local Ethics Committees of IDIBELL-CEEA. Mice were kept in ventilated cages under specific pathogen-free conditions (IDIBELL animal facility). All mice were crossed onto the C57BL/6J genetic background. Alk1+/− mice (Acvrl1+/-) (Oh *et al*, 2000) were a generous gift from Dr. Miguel Pericacho (University of Salamanca, Spain). ALK12loxP;Cdh5(PAC)-CreERT2 mouse line (Sung *et al*, 2009; Ouarné *et al*, 2024) was kindly provided by Dr. Cláudio A. Franco (University of Lisboa, Portugal) as an Acvrl1EC-KO mouse model.

### Pharmacological In Vivo Treatment

For pharmacological experiments, *Acvrl1^flox/flox^*; Cdh5-iCreERT2 and *Acvrl1^flox/flox^* mice were used at the age of 2-month-old. All mice were treated intraperitoneally with 100µL of 10mg/mL tamoxifen for 5 consecutive days. 50% of mice in each group were treated with vehicle solution (0.5% carboxymethyl cellulose) and the other 50% of mice were treated with 37.5mg/kg of selumetinib (MedChemExpress; #HY-50706). Both vehicle and selumetinib were administered via oral gavage in a volume of 200µL in parallel to tamoxifen administration. At the end of the treatment, mice were euthanized by cervical dislocation and necropsy was performed.

### Hematoxylin-Eosin (H&E) Staining

Biopsies were fixed in a formol buffer, dehydrated, and embedded in paraffin. Then, 5µm paraffin-embedded tissue sections were deparaffinized using a battery of 4 xylene trays for 10 minutes each and rehydrated using a battery of graded alcohol for 5 minutes each (three absolute ethanol trays, three 96 % ethanol trays and one 70 % ethanol tray). Afterwards, sections were washed with distilled water and subsequently sunken on hematoxylin for 1 minute. Slides were washed with running tap water and then sunken on 0.5 % HCl solution for 2 seconds and subsequently ammonia solution (1 mL of 30 % ammonia in 200 mL distilled water). Sections were washed with water. Next, they were counterstained with eosin (2.5 g of eosin in 1 L 50% ethanol). Afterwards, sections were dehydrated in graded alcohols followed by xylene. Finally, slides were covered with cover slips and mounted using DPX (Merck).

### Immunohistochemistry

Samples from patients or mice were fixed in 4% buffered formaldehyde, pH 7 and embedded in paraffin. Paraffin-embedded tissue sections were deparaffinized in xylene and rehydrated in downgraded alcohol solutions and distilled water. Antigens were retrieved at pH 6.0 in a Decloaking ChamberTM NxGen (Biocare Medical) under high pressure conditions for 15 minutes at 110 °C. Then endogenous peroxidases were deactivated for 15 minutes with deactivating solution prepared with 3% hydrogen peroxide and 16 % methanol. Samples were subsequently blocked with 2.5% Normal Goat Serum (Vector Laboratories, #30024) for 1 hour. Afterwards, primary antibody (pERK (1:50 Cell Signaling 4376) or Ki67 (1:70 ThermoFisher Scientific MA5-14520)) incubation was performed overnight at 4°C. The day after, samples were washed three times during 10 minutes with 0.1% Triton X-100 1X PBS. Then, tissue sections were incubated for 1 hour at room temperature with secondary antibody IMPRESS HRP Anti-Rabbit IGG PEROX (Vector Laboratories; #480914). Next, samples were washed three times during 10 minutes with 0.1% Triton X-100 1X PBS. Afterwards, staining was revealed by use of the Dako DAB+ Substrate Chromogen developing system (Vector Laboratories, #416425), which was carefully applied on top of each sample for 8 minutes. Finally, samples were counterstained with hematoxylin (Sigma-Aldrich; #517282) for ten minutes. After, samples were washed and subsequently dehydrated in upgraded alcohol solutions followed by xylene in order to be mounted on glass slides with DPX Mountant (Sigma-Aldrich; #06522) mounting medium. Samples were visualized under light microscopy by using a Nikon Eclipse E400 optical microscope.

### Hemorrhage quantifications

Lungs from mice of the different groups were fixed in 4% buffered formaldehyde, pH 7 and embedded in paraffin. Paraffin-embedded tissue sections were desparaffinized in xylene and rehydrated in downgraded alcohol solutions and distilled water. Taking advantage of the hemoglobine peroxidase activity, we detected hemoglobine accumulation in tissues by using Dako DAB+ Substrate Chromogen developing system (Vector Laboratories; #416425), which was applied on top of each sample for 10 minutes. Finally, samples were counterstained with eosin. Subsequently dehydrated in upgraded alcohol solutions followed by xylene, in order to be mounted on glass slides with DPX Mountant (Sigma-Aldrich; #06522) mounting medium. Samples were visualized under light microscopy by using Nikon Elipse E400 optical microscope at 4x magnification. Hemorrhage staining was quantified by Qupath, analysing the percentage of the area affected by hemorrhage relative to the total tissue area.

### Immunofluorescence

Paraffin-embedded sections were deparaffinized using xylene and rehydrated through a graded ethanol series followed by distilled water. Antigen retrieval was conducted at pH 6.0 in a Decloaking Chamber™ NxGen (Biocare Medical) under high-pressure conditions (110°C for 15 minutes). Non-specific binding was blocked by incubating the sections with 5% bovine serum albumin (BSA), 0.1% Donkey Serum, 0.01% Triton in phosphate-buffered saline (PBS) for 1 hour at room temperature. Primary antibodies targeting pERK (1:50 Cell Signaling, 9101S) and ENG (1:100 R&D Systems, AF1097) in the case of human samples, Griffonia Simplicifolia Lectin I (GSL I) isolectin B4 DyLight 594 8 ug/mL (Vector Laboratories, DL-1207-.5) and pERK (1:50 Cell Signaling, 9101S) in mouse samples, were applied and incubated overnight at 4°C. The following day, sections were washed three times for 10 minutes each in PBS containing 0.1% Triton X-100. This was followed by 1-hour incubation at room temperature with h secondary antibodies conjugated to Alexa Fluor dyes. Slides were then washed and stained with DAPI (1:10000) for 10 minutes, and finally mounted using Fluoromount™ mounting medium. Fluorescent imaging was performed using a Nikon Eclipse 80i microscope at 20x or 60x magnification.

### Cell culture

#### Cell Lines

Human umbilical vein endothelial cells (HUVECs) were obtained from Lonza (#C-2517A), and were never used beyond passage eight. hTERT neonatal dermal microvascular endothelial cells (HDMECs) were obtained from ATCC (#CRL-4060). *SMAD6* knockout HDMECs (*SMAD6^KO^* HDMEC) were described in (Cerdà *et al*, 2024). Cells were cultured in 1% gelatin-coated plates with EGM-2 medium (PromoCell) supplemented with 10% FBS, 1% penicillin/streptomycin, 1% L-glutamine and 1% pyruvate; referred as complete EGM. For factor stimulation cells were starved for 16 hours previous to the experiment with DMEM medium (Biowest) containing 0% FBS, 1% penicillin/streptomycin, 1% L-glutamine and 1% pyruvate; referred as depleted medium. Cells were always maintained at 37°C in 5% CO2 atmosphere.

#### siRNA Transfection

100,000 cells were seeded in 1% gelatin-coated 6-well plates and cultured overnight complete EGM2 medium. Next day, cells were transfected with a control siRNA (Thermo Fisher; #4392420) or with the specific siRNA: PPP1R3C (QIAGEN; #1027416-GS5507), SMAD1 (Dharmacon, L-012723-00-0005). To this end, cells were washed with 1X phosphate-buffered saline (PBS) and incubated overnight in 500µL of transfection EGM, containing transfection mix: 100 µL of OptiMEM (Gibco; #31985062), 5µL of 2 µM siRNA, and 1µL of RNAiMAX (Invitrogen; #13778075). Transfection mix was previously incubated for 30 minutes at room temperature. Next day, cells were refreshed with complete EGM2 medium and cultured for additional 48 hours.

### BrdU Incorporation

500,000 cells were seeded in a 12-well plate with coverslip glass on the bottom of each well. Cells were left in the incubator with complete EGM2 medium for 24 hours. 10 µM BrdU (Sigma-Aldrich; #B5002) dissolved in PBS was added into each well and cells were incubated at 37 °C and 5% CO2 for 4 hours. When the treatment was completed, cells were washed thrice with PBS (8 mM Na2HPO4, 2mM KH2HPO4, 154 mM NaCl) and then fixed with 4% PFA in PBS for 15 minutes at room temperature. Cells were washed with PBS. To denature the DNA, cells were treated with 2M HCl for 10 minutes. HCl was neutralized with a borate solution (16.6 mM Na2B4O7 and 83.3mM H2BO3, pH 8.5). Cells were washed once with PBS 0.1% Triton 100X. Samples were blocked with PBS 2% BSA for 40 minutes at room temperature. Next, they were incubated for 3 hours with a rat anti-BrdU-monoclonal antibody (Abcam; #AB6326) diluted 1:250 in blocking buffer. When the incubation was completed, coverslips were washed three times with PBS 0.1 % Triton 100X. Then, samples were incubated for 1 hour with an anti-rat secondary antibody conjugated with Alexa 564, diluted 1:200 in blocking buffer. Subsequently, samples were washed three more times with PBS 0.1% Triton 100X and incubated with 20µg/mL DAPI for 10 minutes. The coverslips were washed once with PBS 0.1% Triton 100, once with PBS and once with distilled H2O. Finally, coverslips were mounted with FluoromountTM Aqueous mounting medium (Sigma; #F4680) and fluorescence was assessed by a fluorescence microscope. Cells were visualized under 20x magnification on a Nikon Eclipse 80i fluorescence microscope, and three images of each coverslip were taken through NIS elements BR software. Counts were performed by using ImageJ software and BrdU incorporation values were obtained through determination of a BrdU-positive nuclei / total nuclei ratio.

### Western Blotting

Cells were washed three times with cold 1X PBS and harvested on ice via cell scraping using 4°C RIPA lysis buffer (1X PBS pH 7.4, 0.1% SDS, 1% NP-40, 0.5% sodium deoxycholate). Lysis buffer was supplemented with a protease and phosphatase inhibitor cocktail (Sigma-Aldrich). Lysates were centrifuged at 16,000g for 15 minutes at 4◦C, and the supernatants were carefully transferred to clean 1.5 mL tubes. Protein concentrations were quantified using the Pierce™ BCA Protein Assay Kit (Thermo Fisher; #23225). Equal amounts of protein from each sample were resolved by SDS-PAGE using the Laemmli buffer system, followed by transfer to Immobilon-P PVDF membranes (Millipore; # IPVH00010) in transfer buffer (25mM Tris/HCl, 190mM glycine, 10% methanol) at 4°C. Membranes were blocked with 5% fat-free powdered milk in 1X TBS (10mM Tris/HCl, 150mM NaCl, pH 7.4) for 1 hour at room temperature and then incubated overnight at 4°C with the appropriate primary antibodies (described in Supplementary Table 1) diluted in 1% fat-free milk in 1X TBS. Following incubation, membranes were washed three times for 10 minutes with 0.1% Triton X-100 in 1X TBS and then probed with horseradish peroxidase-conjugated secondary antibodies—either anti-rabbit IgG (Amersham Pharmacia Biotech; #NA934) or anti-mouse IgG (Amersham Pharmacia Biotech; #NXA931), diluted in 1% fat-free milk in 1X TBS. After additional three washes in 0.1% Triton X-100 in 1X TBS, membranes were incubated for 1 minute with ECL detection reagent, prepared by mixing 1M Tris pH 8.5, 90mM p-coumaric acid, and 250mM luminol in DMSO with 30% hydrogen peroxide at a 1 mL:3µL ratio. Chemiluminescent signals were captured using the ChemiDoc™ MP Imaging System, and band intensity was quantified using ImageLab software (Bio-Rad, Hercules, CA, USA).

### RNA Extraction, Reverse Transcription and qPCR

For RNA extraction, cells cultured in 10 cm plates were lysed in 1 mL of TRIzol™ Reagent (Ambion, Thermo Fisher Scientific; #15596018). For phase separation, 200μL of chloroform were added per 1 mL of TRIzol. Samples were vortexed for 2 minutes and centrifuged at 12,000×g for 15 minutes. The aqueous (upper) phase was carefully collected and mixed 1:1 with isopropanol. Samples were then stored at-20°C for 24 hours to facilitate RNA precipitation, followed by centrifugation at 12,000×g for 30 minutes at 4°C. The supernatant was discarded, and the RNA pellet was washed twice with 1 mL of 70% ethanol. Wash subsequent centrifugation was carried out at 12,000×g for 5 minutes, after which the supernatant was carefully removed. Then, pellets were washed with 1mL of 70% ethanol and resuspended in RNAse free water with 10X Turbo DNAse Buffer and DNase I (ThermoFisher Scientific; Cat. Num. AM1907) to a final volume of 30μL, and incubated for 25 minutes at 4°C. Finally, 3μL of DNAse Inactivation Reagent were added. Then, 2000ng of RNA quantified in a NanoDrop™ spectrophotometer (Thermo Fisher Scientific) were added to a final volume of 20μL and incubated 10 minutes at 65°C. Afterwards, cDNA was obtained by use of a Reverse Transcription Kit (Applied Biosystems, Thermo Fisher Scientific; Cat. Num. #10400745), programmed to be performed in the GeneAmp™ PCR System 9700 Thermal Cycler (ThermoFisher) at 37°C for 2 hours. After that, RT-PCR was performed in a 384-well plate and with SYBR™ Green Master Mix (Applied Biosystems; Cat. Num. A25742). RT-PCR was programmed to be performed up to 40 cycles of amplification in the LightCycler® 480 System (Roche Molecular Biochemicals, Basel, Switzerland). β-Actin was used as a housekeeping gene and CP values were calculated according to the formula: 10000*2-ΔCt. Primers used are listed in Sup. Table 2.

### Bioinformatic Analysis

#### RNA-seq Data Processing

Homo sapiens whole transcriptome sequencing was performed in order to examine the different gene expression profiles, and to perform gene annotation on set of useful genes based on gene ontology pathway information. RNA from control HDMECs, *SMAD6^KO^* HDMEC and *SMAD6^KO^* HDMEC overexpressing WT SMAD6 protein (4 replicates each) was successfully analyzed by Macrogen company. The high-quality trimmed reads were mapped to the reference genome using HISAT2, a highly efficient splice-aware aligner designed to accurately capture exon-exon junctions. Following alignment, transcript assembly was performed with StringTie, which reconstructs full-length transcripts from the aligned reads, enabling a comprehensive representation of transcriptomic structures. Gene expression profiles were then generated, represented both as raw read counts and as normalization values, taking into account transcript length and sequencing depth. Specifically, normalization metrics such as FPKM (Fragments Per Kilobase of transcript per Million mapped reads), RPKM (Reads Per Kilobase of transcript per Million mapped reads), and TPM (Transcripts Per Kilobase Million) were calculated to allow for robust comparisons across samples. Finally, for comparative analyses between the different cell lines, genes or transcripts exhibiting significant differential expression were identified through rigorous statistical hypothesis testing, ensuring a high level of confidence in the observed biological differences.

#### Differentially Expressed Gene Analysis

Statistical analysis of differentially expressed genes (DEG) was performed with Fold Change, nbinomWaldTest, using DESeq2. The significant results were selected on conditions of |fc|>=2 & nbinomWaldTest raw p-value<0.05. Hierarchical clustering analysis (Euclidean Method, Complete Linkage) was used to cluster the similarity of genes and samples by expression level (rlog transformed value) from significant list.

For the enrichment analysis, a test based on the Gene Ontology (GO) database (http://geneontology.org/) was conducted using the list of significantly differentially expressed genes, employing the g:Profiler tool (https://biit.cs.ut.ee/gprofiler/). This tool performs statistical enrichment analysis to identify over-representation of biological information derived from Gene Ontology terms, biological pathways, regulatory DNA elements, human disease gene annotations, and protein-protein interaction networks. The analysis encompassed the three major Gene Ontology categories: biological process, molecular function, and cellular component. Gene products associated with specific GO identifiers were systematically summarized by parsing both the ontology structure file and the annotation file, the latter of which included multispecies annotations provided by UniProt as well as annotations sourced from reference databases associated with the GO Consortium. This approach enabled a comprehensive exploration of the functional enrichment patterns related to the experimental conditions under investigation.

#### RNA-seq Graphical Representation

Heatmaps were created using the website Heatmapper (http://www.heatmapper.ca/expression/), were normalized FPKM values from the different replicates were introduced. The log2 of Fold Change and the-log10 of Benjamini-Hochberg (bh) adjusted p-value were calculated and introduced in GraphPad Prism software for volcano plot representations. Biological processes were filtered by introducing the concept “Cell regulation” in the GO list. The top 10 was selected and represented in a bar graph by introducing the intersection size in GraphPad Prism software.

## Statistical Analysis

Statistical analyses were performed with GraphPad Prism (versions 6.01 and 8.3). Data are mainly presented as Mean ±SD, and each graph dot represents an independent experiment in the case of cells, an independent mouse in the case of adult mice experiments, or an independent blood vessel in the case of patient sample. Different statistical tests were selected based on the distribution characteristics of the data. The normality of data distributions was assessed using the Shapiro-Wilk test for small to moderate sample sizes and the Kolmogorov-Smirnov test for larger sample sizes. Thus, normally distributed data with equal variances were analyzed using the Student’s t-test to compare means between two groups. For non-normally distributed data or with unequal variances, statistical significance was calculated by the non-parametric Mann-Whitney U-test, when comparing two groups. For experiments involving multiple groups, one-way ANOVA was performed to identify overall significant differences.

Statistical significance was determined at a threshold p-value of p<0.05, with specific levels of significance: *p<0.05, **p<0.01, ***p<0.001, ****p<0.0001. All statistical analyses were conducted with rigorous adherence to the assumptions and conditions of each test, ensuring the reliability of the results. In some indicated exceptions, and only if the distribution did not show outliers, normality was assumed without a distribution test due to small sample size.

## Author contributions

Conception and design: JLR, ARM and FV. Acquisition of data (provided animals, acquired and managed patients, provided essential reagents, etc.): JLR, AC, AF, AL, PG, EB, AV, AB-P, MR, JMP, AY, MT-F, AM. Analysis and interpretation of data (e.g., statistical analysis, biostatistics, computational analysis): JLR, PC, ARM, FV. Writing, review, and/or revision of the manuscript: JLR, PC, ARM, FV. Study supervision: ARM, FV.

## Acknowledgements

We particularly wish to acknowledge the collaboration of the patients. We thank Dr. Miguel Pericacho (University of Salamanca, Spain) and Dr. Cláudio A. Franco (University of Lisboa, Portugal) by the generous gift of mouse models.

